# mRNA-1273 vaccine induces neutralizing antibodies against spike mutants from global SARS-CoV-2 variants

**DOI:** 10.1101/2021.01.25.427948

**Authors:** Kai Wu, Anne P. Werner, Juan I. Moliva, Matthew Koch, Angela Choi, Guillaume B. E. Stewart-Jones, Hamilton Bennett, Seyhan Boyoglu-Barnum, Wei Shi, Barney S. Graham, Andrea Carfi, Kizzmekia S. Corbett, Robert A. Seder, Darin K. Edwards

## Abstract

Severe acute respiratory syndrome coronavirus-2 (SARS-CoV-2) is the causative infection of a global pandemic that has led to more than 2 million deaths worldwide. The Moderna mRNA-1273 vaccine has demonstrated ~94% efficacy in a Phase 3 study and has been approved under Emergency Use Authorization. The emergence of SARS-CoV-2 variants with mutations in the spike protein, most recently circulating isolates from the United Kingdom (B.1.1.7) and Republic of South Africa (B.1.351), has led to lower neutralization from convalescent serum by pseudovirus neutralization (PsVN) assays and resistance to certain monoclonal antibodies. Here, using two orthogonal VSV and lentivirus PsVN assays expressing spike variants of 20E (EU1), 20A.EU2, D614G-N439, mink cluster 5, B.1.1.7, and B.1.351 variants, we assessed the neutralizing capacity of sera from human subjects or non-human primates (NHPs) that received mRNA-1273. No significant impact on neutralization against the B.1.1.7 variant was detected in either case, however reduced neutralization was measured against the mutations present in B.1.351. Geometric mean titer (GMT) of human sera from clinical trial participants in VSV PsVN assay using D614G spike was 1/1852. VSV pseudoviruses with spike containing K417N-E484K-N501Y-D614G and full B.1.351 mutations resulted in 2.7 and 6.4-fold GMT reduction, respectively, when compared to the D614G VSV pseudovirus. Importantly, the VSV PsVN GMT of these human sera to the full B.1.351 spike variant was still 1/290, with all evaluated sera able to fully neutralize. Similarly, sera from NHPs immunized with 30 or 100μg of mRNA-1273 had VSV PsVN GMTs of ~ 1/323 or 1/404, respectively, against the full B.1.351 spike variant with a ~ 5 to 10-fold reduction compared to D614G. Individual mutations that are characteristic of the B.1.1.7 and B.1.351 variants had a similar impact on neutralization when tested in VSV or in lentivirus PsVN assays. Despite the observed decreases, the GMT of VSV PsVN titers in human vaccinee sera against the B.1.351 variant remained at ~1/300. Taken together these data demonstrate reduced but still significant neutralization against the full B.1.351 variant following mRNA-1273 vaccination.

## INTRODUCTION

Moderna’s SARS-CoV-2 vaccine, mRNA-1273, elicits high viral neutralizing titers in Phase 1 trial participants (Jackson et al, 2020; Anderson et al, 2020) and is highly efficacious in prevention of symptomatic COVID-19 disease and severe disease (Baden et al., 2020). However, the recent emergence of SARS-CoV-2 variants in the United Kingdom (B.1.1.7 lineage) and in South Africa (B.1.351 lineage) has raised concerns due to their increased rates of transmission as well as their potential to circumvent immunity elicited by natural infection or vaccination (Volz et al., 2021; Tegally et al., 2020; Wibmer et al., 2021; Wang et al., 2021; Collier et al., 2021).

First detected in September 2020 in South England, the SARS-CoV-2 B.1.1.7 variant has spread at a rapid rate and is associated with increased transmission and higher viral burden (Rambaut et al., 2020). This variant has seventeen mutations in the viral genome. Among them, eight mutations are located in the spike (S) protein, including 69-70 del, Y144 del, N501Y, A570D, P681H, T716I, S982A and D1118H. Two key features of this variant, the 69-70 deletion and the N501Y mutation in S protein, have generated concern among scientists and policy makers in the UK based on increased transmission and potentially increased mortality, resulting in further shutdowns. The 69-70 deletion is associated with reduced sensitivity to neutralization by SARS-CoV-2 human convalescent serum samples (Kemp et al, 2021). N501 is one of the six key amino acids interacting with ACE-2 receptor (Starr et al. 2020), and the tyrosine substitution has been shown to have increased binding affinity to the ACE-2 receptor (Chan et al., 2020).

The B.1.351 variant emerged in South Africa over the past few months, and, similar to the B.1.1.7 variant, increased rates of transmission and higher viral burden after infection have been reported (Tegally et al., 2020). The mutations located in the S protein are more extensive than the B.1.1.7 variant with changes of L18F, D80A, D215G, L242-244del, R246I, K417N, E484K, N501Y, D614G, and A701V, with three of these mutations located in the RBD (K417N, E484K, N501Y). B.1.351 shares key mutations in the RBD with a reported variant in Brazil (Tegally et al., 2020; Naveca et al., 2021). As the RBD is the predominant target for neutralizing antibodies, these mutations could impact the effectiveness of monoclonal antibodies already approved and in advanced development as well as of polyclonal antibody elicited by infection or vaccination in neutralizing the virus (Greaney et al., 2021, Wibmer et al, 2021).

Recent data have suggested that the key mutation present in the B.1.351 variant, E484K, confers resistance to SARS-CoV-2 neutralizing antibodies, potentially limiting the therapeutic effectiveness of monoclonal antibody therapies (Wang et al., 2021; Greaney et al., 2020; Weisblum et al., 2020; Liu et al., 2020; Wibmer et al., 2021). Moreover, the E484K mutation was shown to reduce neutralization against a panel of convalescent sera (Weisblum et al., 2020; Liu et al., 2020; Wibmer et al., 2021). In terms of vaccination, it is clear that mRNA-1273 induces significantly higher neutralizing titers than convalescent sera against the USA-WA1/2020 isolate (Jackson et al, 2020). A recent study using a recombinant VSV PsVN assay showed that sera of mRNA-1273 vaccinated participants had reduced neutralizing titers against E484K or K417N/E484K/N501Y combination (Wang et al, 2021), however there has been no assessment of sera from mRNA-1273 clinical trial participants against the full constellation of S mutations found in the B.1.1.7 or B.1.351 variants.

In this study, we assessed neutralization of sera from mRNA-1273 vaccinated Phase 1 clinical trial participants against recombinant VSV-based SARS-CoV-2 PsVN assay with S protein from the original Wuhan-Hu-1 isolate, D614G variant, the B.1.1.7 and B.1.351 variants, and variants that have previously emerged (20E, 20A.EU2, D614G-N439K, and mink cluster 5 variant). We also assessed the effect of both single mutations and combinations of mutations present in the RBD region of the S protein. In addition, orthogonal assessments in VSV and pseudotyped lentiviral neutralization assays were performed on sera from NHPs that received the mRNA-1273 vaccine at two different dose levels, as this has been a useful pre-clinical model for vaccine induced immunogenicity and protection. Using both of these assays provided confirmatory data on pseudovirus neutralization. Overall, this comprehensive pseudovirus neutralization analysis in humans and non-human primates that received mRNA-1273 provides critical initial data necessary to elucidate how vaccines may be impacted by SARS-CoV-2 variants.

## RESULTS

### mRNA-1273 neutralization against mutant strains

To assess the ability of mRNA-1273 to elicit neutralizing antibodies against the new SARS-CoV-2 variants, we first evaluated sera from NHPs that received 30 μg of mRNA-1273 and participants in the Phase 1 clinical study immunized with mRNA-1273 at the authorized dose of a 100 μg; both NHPs and humans received a primary series - two doses given approximately 28 days apart. Neutralizing activity was measured with SARS-CoV-2 full-length S pseudotyped recombinant VSV-ΔG-firefly luciferase virus; antibody levels were measured in PsVN assays containing the S of the original Wuhan isolate (D614), the dominant strain of 2020 containing the D614G mutation, or S from 20E, 20A.EU2 and mink cluster 5 variants. (**Table 1**). Results demonstrate that the antibody response elicited by mRNA-1273 provides similar levels of neutralization against these SARS-CoV-2 S variants as against the Wuhan-Hu-1 (D614) strain. This observation includes the G614 variant that has been shown to have higher neutralizing titers in lentiviral PsVN assays **(Fig. 1)**.

**Table 1.** Spike mutations in SARS-CoV-2 variants evaluated in this study.

| Variant Name | Amino Acid Changes in Spike |
| --- | --- |
| **20E (EU1)** | A222V-D614G |
| **20A.EU2** | S477N-D614G |
| **N439K-D614G** | N439K-D614G |
| **Mink Cluster 5 Variant** | ΔH69ΔV70-Y453F-D614G-I692V-M1229I |
| **B.1.1.7**<br>(a.k.a., 20I/501Y.V1, VOC 202012/01) | ΔH69ΔV70-ΔY144-N501Y-A570D-D614G-P681H-T716I-S982A-D1118H |
| **B.1.351**<br>(a.k.a., 20H/501Y.V2) | L18F-D80A-D215G-ΔL242ΔA243ΔL244-R246I-K417N-E484K-N501Y-D614G-A701V |

**Figure 1.**
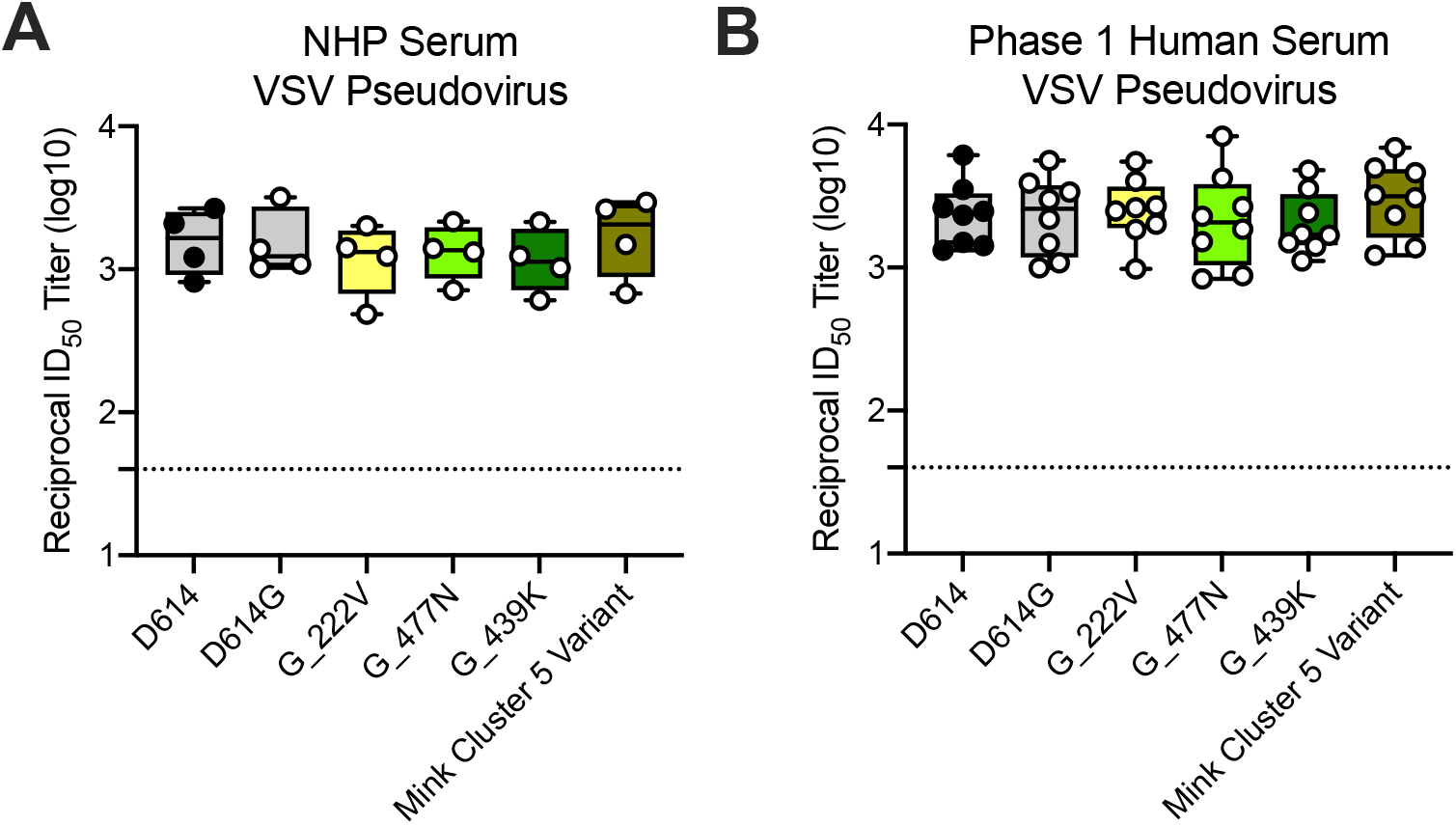
Ability of mRNA-1273 immune sera from NHPs and humans to neutralize SARS-CoV-2 pseudoviruses representing early variants. **(A)** Rhesus macaques (NHPs) were immunized with 30 μg mRNA-1273 on a prime-boost schedule, and sera were collected 4 weeks post-boost. **(B)** Phase 1 trial participants were immunized with 100 μg mRNA-1273 on a prime-boost schedule, and sera were collected 1 week post-boost. Neutralization was measured by a recombinant VSV-based SARS-CoV-2 pseudovirus neutralization assay incorporating full-length spike protein of the Wuhan isolate (D614) or the indicated spike variants (D614G, A222V-D614G, S477N-D614G, N439K-D614G, mink cluster 5 variant). Min to max box plots, with the box from 25-75% and the median value denoted by the line. The horizonal dotted lines indicate the lower limit of quantification (LLOQ=40). G=D614G.

### mRNA-1273 NHP neutralization against B. 1.1.7 and B. 1.351

We next evaluated sera from NHPs vaccinated with a primary series of 30 or 100 μg mRNA-1273. Neutralizing antibody responses were measured using both VSV - based PsVN assay and our standardly-reported luciferase-based lentiviral assay. Pseudoviruses in both assays incorporate the full-length S, encoding the original Wuhan-Hu-1 (D614), D614G, single, partial or complete set of mutations that are present in B.1.1.7 and B.1.351 lineages **(Table 2)**.

**Table 2.** Spike variants evaluated in PsVN assay assessment of mRNA-1273 vaccinated NHP sera

| Pseudovirus | Strain | Partial or full Spike mutants | Mutations |
| --- | --- | --- | --- |
| Lentivirus | Control |  | D614G |
|  | B.1.1.7 & B.1.351 | Partial | D614G-N501Y |
|  | B.1.1.7 | Partial | D614G-N501Y-ΔH69ΔV70 |
|  | B.1.351 | Partial | D614G-E484K |
|  | B.1.351 | Partial | D614G-N501Y-E484K-K417N |
| VSV | Control |  | D614G |
| | B.1.1.7 | Full | ΔH69ΔV70-ΔY144-N501Y-A570D-D614G-P681H-T716I-S982A-D1118H |
|  | B.1.351 | Partial | K417N-E484K-N501Y-D614G |
| | B.1.351 | Full | L18F-D80A-D215G-ΔL242ΔA243ΔL244-R246I-K417N-E484K-N501Y-D614G-A701V |

The mutations present in the B.1.1.7 variant, either the complete set of S mutations or the specific mutations (N501Y, ΔH69ΔV70) of key interest **(Table 2)** had minimal effect on neutralization in both the VSV and lentiviral neutralization assays. In the VSV assay, no difference was observed between the D614 and G614 viruses, and there was potent neutralization measured against both **(Figure 2A)**.

**Figure 2.**
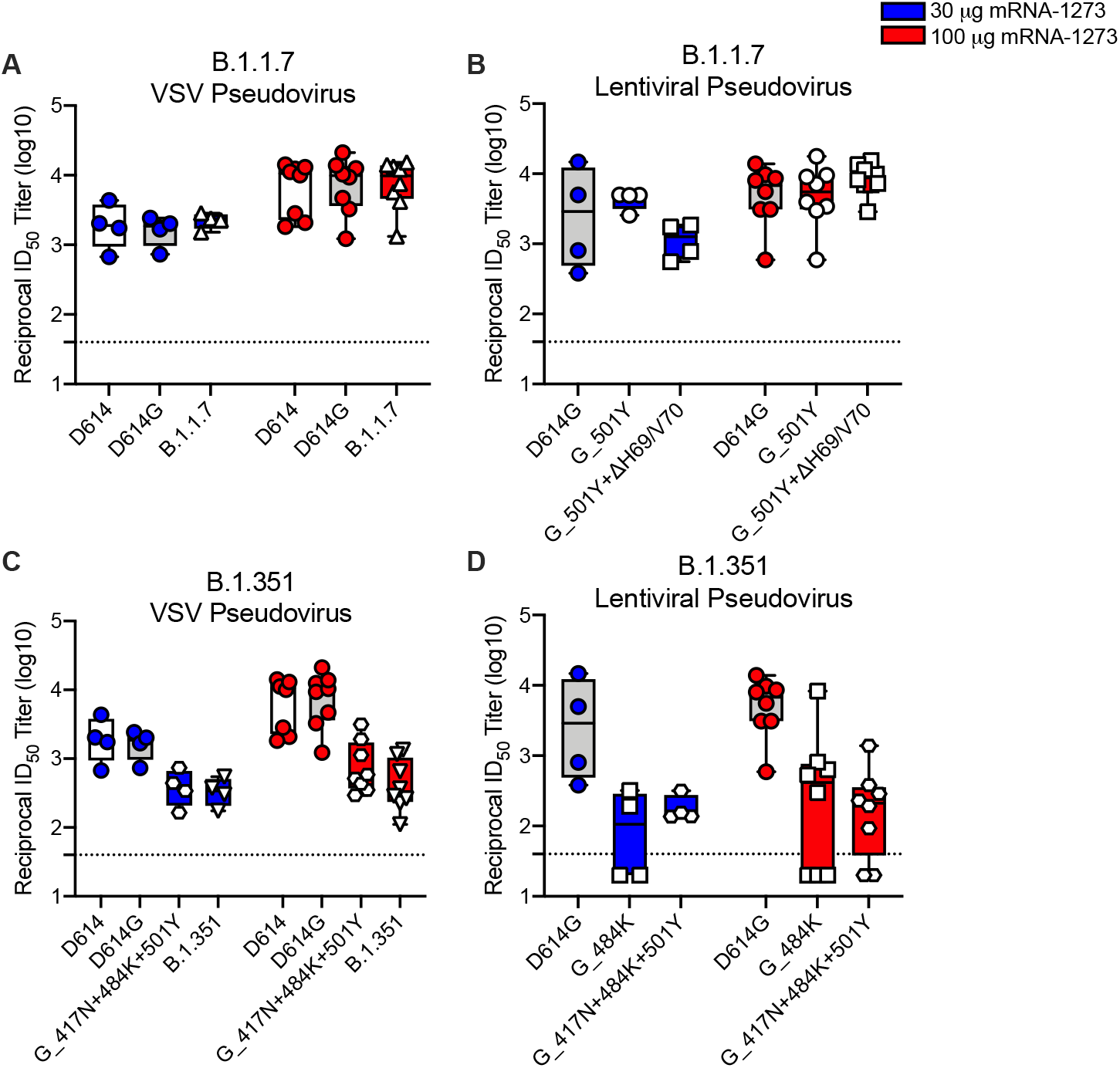
Neutralization of B.1.1.7 and B.1.351 SARS-CoV-2 pseudoviruses by serum from mRNA-1273-immunized NHPs. Rhesus macaques (NHPs) were immunized with 30 (blue) or 100 μg (red) mRNA-1273 on a prime-boost schedule, and sera were collected 4 weeks post-boost. Neutralization was measured by recombinant VSV-based pseudovirus neutralization assay (**A, C**) or lentiviral pseudovirus neutralization assay (**B, D**). The assays incorporated full-length Spike protein of the original D614, D614G, or the indicated Spike variants present in the B.1.1.7 variant (**A, B**) or B.1.351 variant (**C, D**). Min to max box plots, with the box from 25-75% and the median value denoted by the line. The horizonal dotted lines indicate the lower limit of quantification (LLOQ). G=D614G

In the lentiviral assay, a slight drop in neutralizing titers was measured between the G614 (G) and the combined N501Y and H69/V70 deletions. However, despite a 2-fold reduction in the geometric mean ID_50_ titers against G_501Y+ΔH69/V70 pseudotyped lentivirus for NHP receiving 30 μg of mRNA-1273, neutralizing titers remained high and this modest decrease was not reflected in the VSV assay. Additionally, NHPs immunized with the 100 μg dose demonstrated robust neutralizing antibody responses against G_501Y+ΔH69/V70 that were similar to the responses against the G614 virus **(Figure 2B)**.

A significant decrease in neutralizing titers was measured against both the total combined B.1.351 S mutations, and against isolated single mutations or combined mutations found within the B.1.351 variant, as described in **Table 2**. In the VSV assay, a 4.3- and 4.8-fold drop in neutralizing titers from sera collected from 30 μg dosed animals and a 9.6 and >10-fold drop in neutralizing titers from sera from 100 μg dosed animals were measured against the partial or complete B.1.351 mutant pseudovirus, respectively **(Figure 2B)**. All samples were able to fully neutralize the virus, although at lower dilutions as shown by the neutralization curves from the assay **(Figure 3)**. In the lentiviral pseudovirus neutralization assay, the full set of mutations present in the B.1.351 variant were not assessed; instead only key mutations present in the RBD of the S protein were incorporated; namely D614G_E484K or D614G_N501Y_E484K_K417N **(Table 2)**. With this pseudovirus assay, there was a 5.2-fold decrease in geometric mean neutralizing ID50 titers against the single 484K mutant for NHPs immunized with 100 μg mRNA-1273. In some animals in each of the dose groups the geometric mean neutralizing ID50 titers were below the limit of quantification for the assay **(Figure 2D)**.

**Figure 3.**
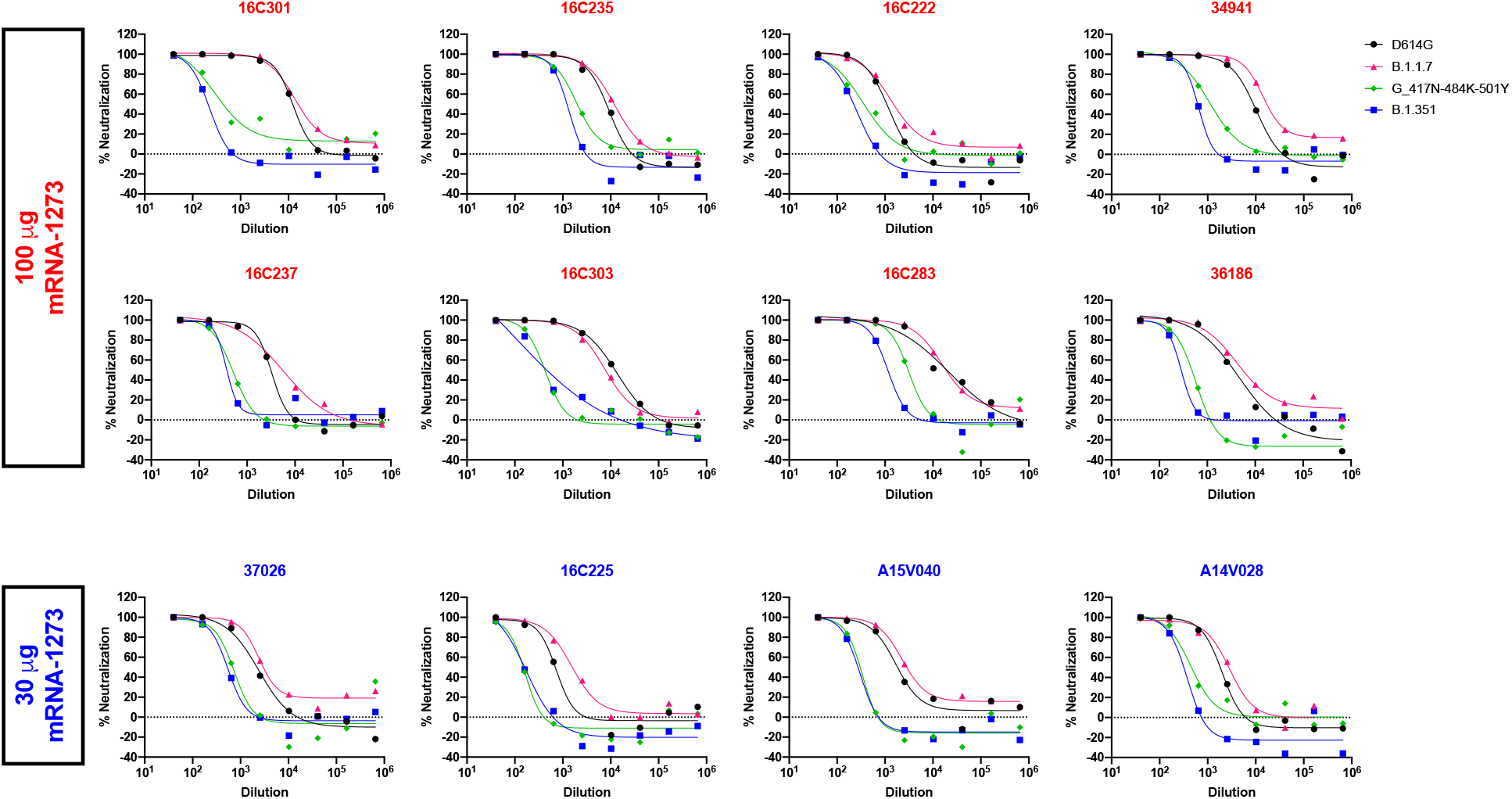
Neutralization curves of NHP sera in the VSV-based pseudovirus neutralization assay. Rhesus macaques (NHPs) were immunized with 30 (blue) or 100 μg (red) mRNA-1273 on a prime-boost schedule, and sera were collected 4 weeks post-boost. Neutralization was measured against recombinant VSV-based pseudovirus incorporating the B.1.351 variant full-length spike protein. Each graph represents an individual animal as indicated by identification code. Each data point is an average from three replicate wells.

### mRNA-1273 Phase 1 human sera neutralization against B. 1.1.7 and B. 1.351

The mutations present in the B.1.1.7 variant, either the full panel of S mutations or key mutations in the RBD region **(Table 2)**, had minimal effect on neutralization of mRNA-1273 Phase 1 participant sera **(Figure 4A-B)**. In contrast, a significant decrease in neutralizing titers was measured against both the full set of S mutations and the partial list of RBD mutations in the B.1.351 variant, listed in **Table 2**. In the VSV assay, using Phase 1 one-week post-boost sera samples, we detected a 2.7- and 6.4-fold reduction in neutralizing titers against the partial or full panel of mutations, respectively **(Figure 4C-D)**. Despite diminished neutralizing responses against the B.1.351 variant, neutralizing titers remain generally high (1/290), and all sera samples completely neutralized the VSV pseudovirus, albeit at lower dilutions as depicted by the neutralization assay curves **(Figure 5)**.

**Figure 4.**
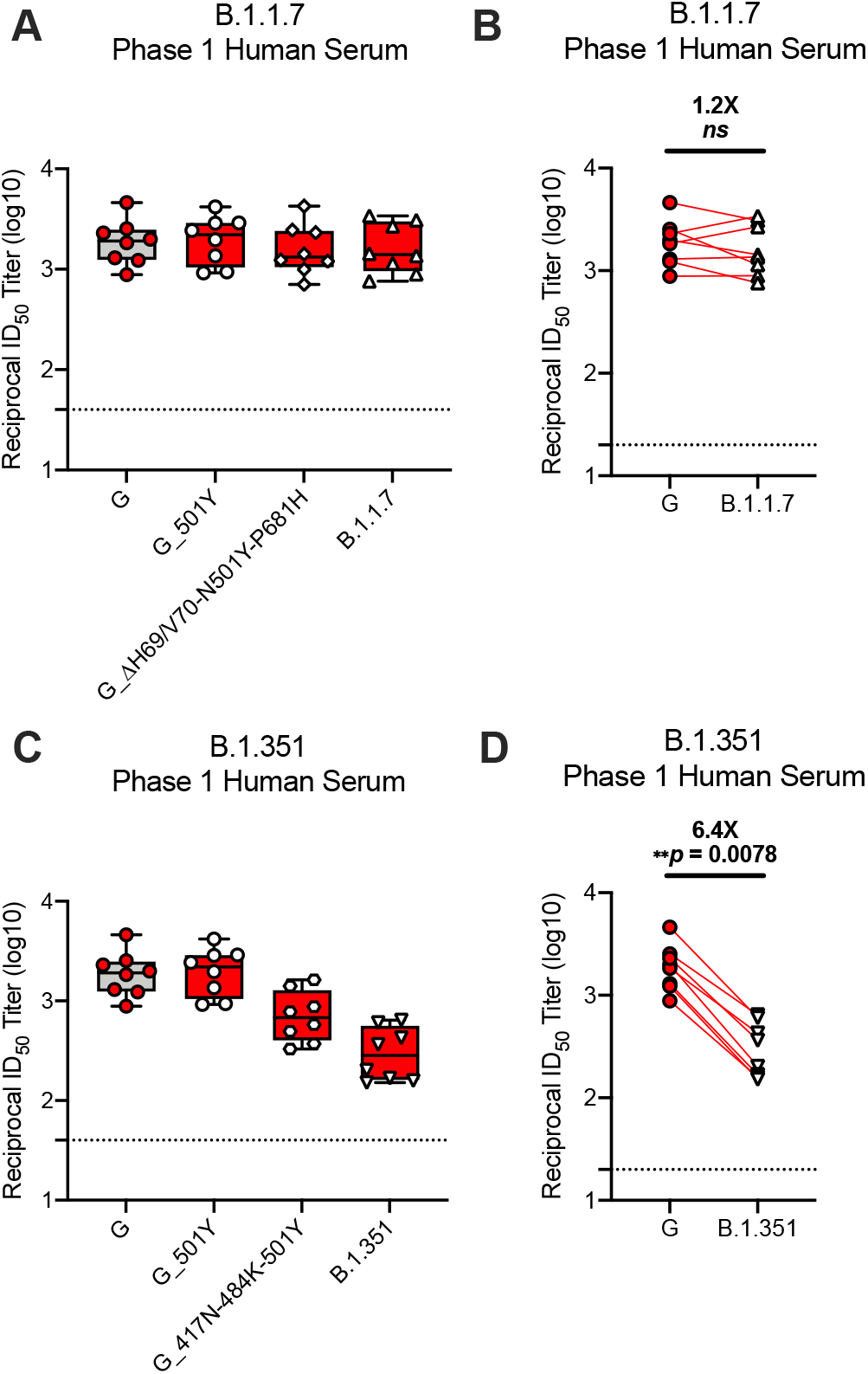
Neutralization of B.1.1.7 and B.1.351 SARS-CoV-2 pseudoviruses by serum from mRNA-1273-immunized Phase 1 participants. mRNA-1273 Phase 1 trial participant sera were collected on day 36, 7 days after the boost. Neutralization was measured by a recombinant VSV-based PsVN assay that incorporated D614G or the indicated spike mutations present in the B.1.1.7 variant (**A-B**) or B.1.351 variant (**C-D**). (**A,C**) Min to max box plots, with the box from 25-75% and the median value denoted by the line. (**B,D**) Results from individual participant sera is represented as dots on each figure, with lines connecting the D614G and variant neutralization titers. The horizonal dotted lines indicate the lower limit of quantification (LLOQ). G=D614G

**Figure 5.**
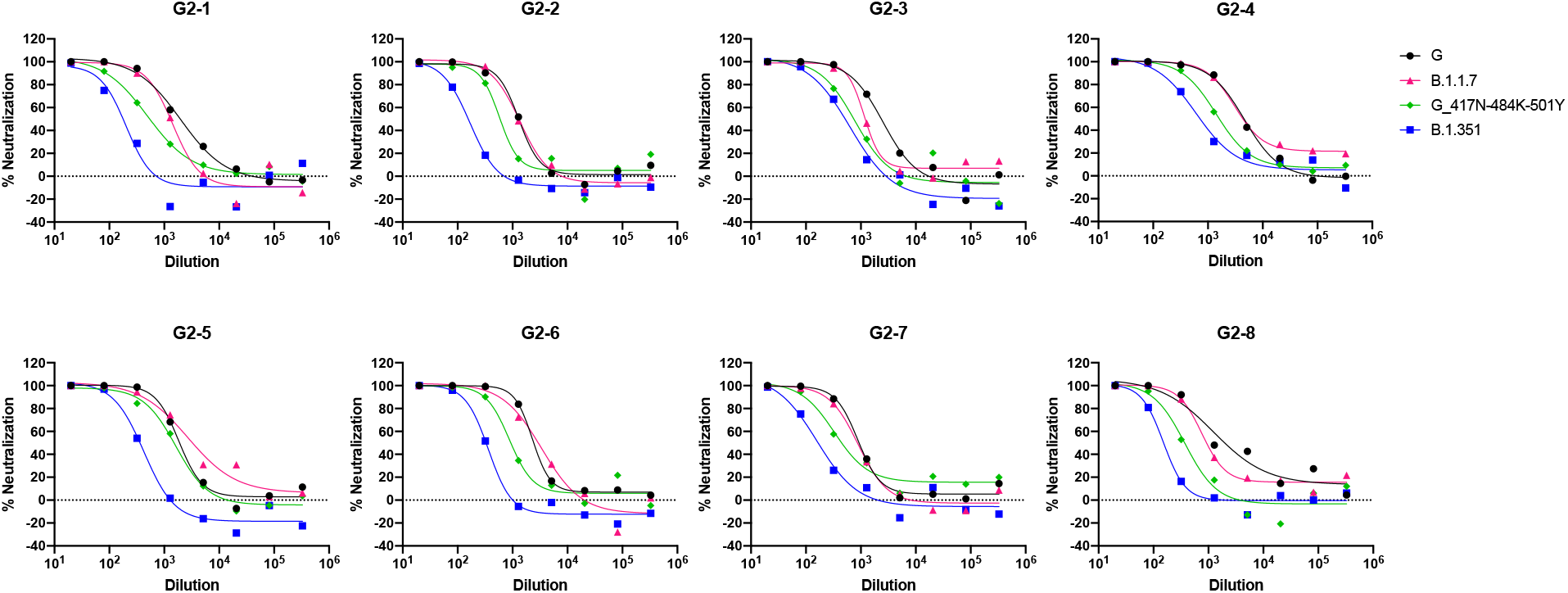
Neutralization curves of human sera in the VSV-based pseudovirus neutralization assay. Phase 1 participants were immunized with 100 μg mRNA-1273 on a prime-boost schedule, and sera were collected 1 week post-boost. Neutralization was measured against recombinant VSV-based pseudovirus incorporating the B.1.351 variant full-length spike protein. Each graph represents an individual participant, as indicated. Each data point is an average from three replicate wells.

## DISCUSSION

We assessed the neutralization capacity of sera from eight phase 1 clinical trial participants (aged 18-55 years) who received two 100 μg doses of mRNA-1273, and NHPs immunized with two doses of 30 μg or 100 μg of mRNA-1273. Neutralization was measured against the original D614 S, the dominant G614 S, mutations in 20E, 20A.EU2, mink cluster 5, N439K-D614G, and either the full panel or single and subset of mutations found in the B.1.1.7 and B.1.351 variants strains. Notably, the 30 μg dose in NHPs elicits similar neutralizing titers against both D614 and G614 VSV pseudoviruses to those of humans receiving two 100 μg doses of mRNA-1273. Assessing the 30 μg dose in NHPs in ongoing and future studies may also help elucidate any dose-dependent effects on neutralizing responses and protection toward the new S variant viruses.

We evaluated both single and combined mutations of interest found in the B.1.1.7 or B.1.351 variants in vitro utilizing both a pseudotyped lentiviral reporter system and a VSV-pseudovirus reporter system. In analyses of both human and NHP sera, we first determined the neutralizing responses against D614 and G614 S to provide a baseline for comparison with the newer variants. Consistent with prior analyses, all eight samples from Phase 1 participants demonstrated robust neutralizing responses against both D614 and G614 S SARS-CoV-2 (Jackson et al., 2020). Additionally, mRNA-1273-immunized NHPs at two different dose levels showed neutralizing antibody titers consistent with established protective efficacy against the WA strain (Corbett et al., 2020).

No significant impact on neutralization was observed from either the full set of mutations found in the B.1.1.7 variant or the N501Y and the 69-70 deletion. Although these mutations have been reported to lessen neutralization from convalescent sera, sera from the Phase 1 participants and NHPs immunized with mRNA-1273 were able to neutralize the B.1.1.7 variant to the same level as the D614G virus.

Consistent with other recent reports assessing neutralization of the mutations found in B.1.351 (Wang et al, 2021), there was a 2.7-fold reduction in neutralization from sera collected from participants vaccinated with mRNA-1273 when the 3 mutations found in the RBD (K417N-E484K-N501Y) were present in the VSV-based pseudovirus assay. Importantly a 6.4-fold reduction was observed when the full set of mutations, including those in the N-terminal domain (NTD), were included. The data in NHP showed a >10 or 4.8-fold reduction from 100 and 30 μg dose groups, respectively, compared to G614. All samples from both the clinical trial participants and NHPs fully neutralized the variant viruses in the VSV PsVN assay, albeit at lower dilutions of sera, and the neutralizing titers remained at ~ 1/300. While prior studies have shown that pseudovirus neutralizing titers against G614 correlate with viral neutralizing titers, it will be important to substantiate these results by testing the B.1.351 variant in a live virus neutralization assay.

Data from this sample set shows mRNA-1273 maintained activity against all circulating strain variants tested to date, and only the B.1.351 variant showed reduced neutralizing titers, as assessed from vaccinated human and NHP sera. Viral escape was not detected from any sample and neutralizing titers remained above those previously found to be protective in NHP challenge studies.

The emergence of strain variants and the ability of the virus to partially overcome natural or vaccine-induced immunity does serve as a call to action, pointing to the need for continued efforts to vaccinate with the currently approved mRNA regimens to prevent the emergence of future variants that may further evade immunity. In addition, active viral surveillance and testing of protection against new viral variants must continue, and if warranted new vaccine efforts must be engaged to protect against breakthrough strains. The mRNA platform allows rapid design of vaccine antigens that incorporate key mutations. Strain-matched vaccines can be developed in response to these variants, either to understand how we might evolve the vaccine-induced immune response through boosting, or to assess the cross-protection provided by a primary series.

The vaccine development community has been mobilized against SARS-CoV-2 and monitoring and vaccine efforts have already shown to be rapid and effective as evidenced by the fact that multiple vaccines were authorized in less than 12 months from the identification of SARS-CoV-2. Future responses will need to maintain this paradigm of rapid translation of the emerging scientific and clinical data into effective medical countermeasures.

## METHODS

### Animal Studies

Experiments in animals were performed in compliance with National Institutes of Health (NIH) regulations and with approval from the Animal Care and Use Committee of the Vaccine Research Center. Female and male Indian-origin rhesus macaques (12 of each sex; age range, 3 to 6 years) were sorted according to sex, age, and weight, and then stratified into groups. Animals were vaccinated intramuscularly at week 0 and at week 4 with either 30 or 100 μg of mRNA-1273 in 1 ml of 1× phosphate-buffered saline (PBS) into the right hind leg. At week 8 (4 weeks after the second vaccination), sera were collected for immunoassay analyses.

### Clinical trial

Humans were immunized with 100 μg mRNA-1273 on a prime-boost schedule and sera was collected 1 week post the boost (day 36). Study protocols and results are reported in Jackson et al, 2020.

### Lentiviral-based Pseudovirus Neutralization

To produce SARS-CoV-2 pseudotyped lentivirus, a codon-optimized CMV/R-SARS-CoV-2 S (Wuhan-1, GenBank: MN908947.3) plasmid was constructed and subsequently modified via site-directed mutagenesis to contain the D614G mutation. Additional spike mutations were implemented into the D614G backbone (i.e. N501Y, E484K, N439K, and other combinations akin to those of the B.1.1.7 and B.1.351 variants). Pseudoviruses were produced by the co-transfection of plasmids encoding a luciferase reporter, lentivirus backbone, and the SARS-CoV-2 S genes into HEK293T/17 cells (ATCC CRL-11268), as previously described (Wang et al., 2015). Additionally, a human transmembrane protease serine 2 (TMPRSS2) plasmid was co-transfected to produce pseudovirus (Böttcher et al., 2006). Neutralizing antibody responses in sera were assessed by PsVN assay, as previously described (Jackson et al., 2020). Briefly, heat-inactivated serum was serially diluted in duplicate, mixed with pseudovirus, and incubated at 37°C and 5% CO2 for roughly 45 minutes. 293T-hACE2.mF cells were diluted to a concentration of 7.5 x 10^4^ cells/mL in DMEM (Gibco) supplemented with 10% Fetal Bovine Serum (FBS) and 1% Penicillin/Streptomycin and added to the serum-pseudovirus mixtures. Seventy-two hours later, cells were lysed and luciferase activity (in relative light units (RLU)) was measured. Percent neutralization was normalized considering uninfected cells as 100% neutralization and cells infected with pseudovirus alone as 0% neutralization. IC_50_ titers were determined using a log(agonist) vs. normalized-response (variable slope) nonlinear regression model in Prism v8 (GraphPad).

### Recombinant VSV-based Pseudovirus Neutralization

Codon-optimized full-length spike protein of the original Wuhan isolate (D614), D614G, or the indicated spike variants listed in Tables 1 and 2 were cloned into pCAGGS vector. To make SARS-CoV-2 full-length spike pseudotyped recombinant VSV-ΔG-firefly luciferase virus, BHK-21/WI-2 cells (Kerafast, EH1011) were transfected with the spike expression plasmid and subsequently infected with VSVΔG-firefly-luciferase as previously described (Whitt, 2010). For neutralization assay, serially diluted serum samples were mixed with pseudovirus and incubated at 37 Celsius for 45 minutes. The virus-serum mix was subsequently used to infect A549-hACE2-TMPRSS2 cells for 18 hr at 37 Celsius before adding ONE-Glo reagent (Promega E6120) for measurement of luciferase signal (relative luminescence unit; RLU). The percentage of neutralization is calculated based on RLU of the virus only control, and subsequently analyzed using four-parameter logistic curve (Prism 8).

## AUTHOR CONTRIBUTIONS

Conceptualization, K.W., A.P.W., B.S.G., A.C., K.S.C., R.A.S, and D.K.E.; methodology, K.W., A. P.W., S.B.B, W.S., and K.S.C.; formal & statistical analysis, K.W. and A.P.W.; writing—original draft preparation, R.A.S. and D.K.E; writing—review and editing, K.W., A.P.W., G.S.J., H.B., B. S.G., A.C., K.S.C., R.A.S., and D.K.E. All authors have read and agreed to the published version of the manuscript.

## ACKNOWLEDGEMENTS

We thank the mRNA-1273 phase 1 study team and the Division of Microbiology and Infectious Diseases, NIAID for providing clinical samples; Dr. Michael Whitt for kind support on recombinant VSV-based SARS-CoV-2 pseudovirus production; Gabriela Alvarado for project management support; members of the VRC Translational Research Program for technical and administrative assistance with NHP experiments; Huihui Mu and Michael Farzan for providing angiotensin-converting enzyme 2 (ACE2)–overexpressing 293 cells; Nicole Doria-Rose for advice regarding lenti-based pseudovirus neutralization assays.

